# Predicting and Designing Red Fluorescent Protein Variants Using Sequence-to-Function Machine Learning Models

**DOI:** 10.64898/2025.12.05.690496

**Authors:** Ran Ji, Jean Jung, Howard Cheng, Ella Y. Xu, Audrey Wang, Keith Pardee, Yufeng Zhao

## Abstract

Fluorescent proteins (FPs) are widely used reporters for visualizing cellular structures and processes. Traditional wet-lab strategies for FP engineering (rational design and directed evolution) have enabled substantial improvements in photophysical performance but are limited by their requirement for deep expert knowledge or labor-intensive screening. AI-driven approaches have recently gained traction for engineering variants of green FPs, yet applications to red fluorescent proteins (RFPs) remain scarce. Here, we applied machine learning models to an RFP sequence-function dataset and trained these models to predict functional single-mutation variants of the state-of-the-art RFP mScarlet-I3. Guided by model predictions, we identified variants exhibiting red-shifted emission peaks, large Stokes shifts, or brightness comparable to the parental protein. Our findings show that even lightweight, data-efficient models can extract actionable design principles for improving RFPs. This work demonstrates the feasibility of AI-guided design for RFPs and provides a reliable benchmark for future development of more powerful AI-driven strategies for FP engineering.

## Introduction

Fluorescent proteins (FPs) have become indispensable tools in biological research, enabling visualization of cellular structures and dynamics with high spatiotemporal resolution1,2. Originally discovered in nature, these proteins have since been extensively engineered to achieve enhanced brightness^3^, expanded spectral diversity^4^, and specialized photophysical properties such as photoswitching^5,6^ and photoactivation^7,8^. Experimental approaches for engineering FP include rational and semi-rational design, which leverage physical and chemical principles to introduce desired interactions between the autocatalytically formed chromophore (from a tripeptide)^9,10^ and its surrounding amino acids. Directed evolution serves as a complementary strategy, relying on screening of mutant libraries generated by random mutagenesis without requiring prior structural knowledge^11^.

Despite many successes, both strategies face inherent limitations. Knowledge-or rule-based design often suffers from low success rates due to the complexity of protein systems. Directed evolution, while powerful, is constrained by practical throughput. Screening libraries of thousands to millions of variants covers only a tiny fraction of the vast protein sequence landscape. For example, the combinatorial possibilities at just six amino acid positions already exceed 60 million variants.

AI-driven approaches to protein engineering are attracting growing interest^12,13^, particularly in light of the landmark successes of structure prediction methods like AlphaFold^14,15^, RoseTTAFold^16^, and others^17^. AI models have been applied to engineer diverse proteins, including channelrhodopsins^18^, enzymes^19,20^, binders^21,22^, and biosensors^23,24^. Green FPs (GFPs) have served as model systems to benchmark AI models, particularly for predicting variants with enhanced brightness^25,26^. Combining structural modeling and energy calculations with AI has proven effective for designing GFP mutants with high quantum yield^27^. In practical applications, AI-guided approaches have also been used to improve state-of-the-art biosensors, such as GFP-based Ca^2+^ indicators with enhanced sensitivity^24^ and sensors re-engineered to alter binding specificity from acetylcholine to serotonin^23^.

In addition to the broadly used GFPs, red FPs (RFPs) offer an orthogonal spectral channel for multiplexed detection, along with the advantages of reduced autofluorescence, scattering, and absorption in biological tissues^28^. However, applications of AI to RFPs remain limited. One possible reason is that models trained on GFPs often fail to generalize well to RFPs, due to the substantial sequence divergence and differences in chromophore structure between the two classes, despite their shared barrel-like protein 3D structure.

In this study, we collected over 150 reported RFP sequences and applied machine learning (ML) strategies to analyze and predict RFP mutants with desirable photophysical properties (**Fig. 1a**). We selected classical ML models with amino acid descriptors for their simplicity and cost-effectiveness. Unlike prior GFP-focused studies that often began with low-to-moderate performing templates (e.g., avGFP or EGFP), we chose the state-of-the-art RFP mScarlet-I3^29^ as our starting point. From a practical standpoint, beginning with the best available template allows us to explore improvements that are directly relevant for advancing FP applications. We further validated the ML predictions experimentally and identified several new mScarlet-I3 variants exhibiting the properties predicted by the model.

**Fig. 1.**
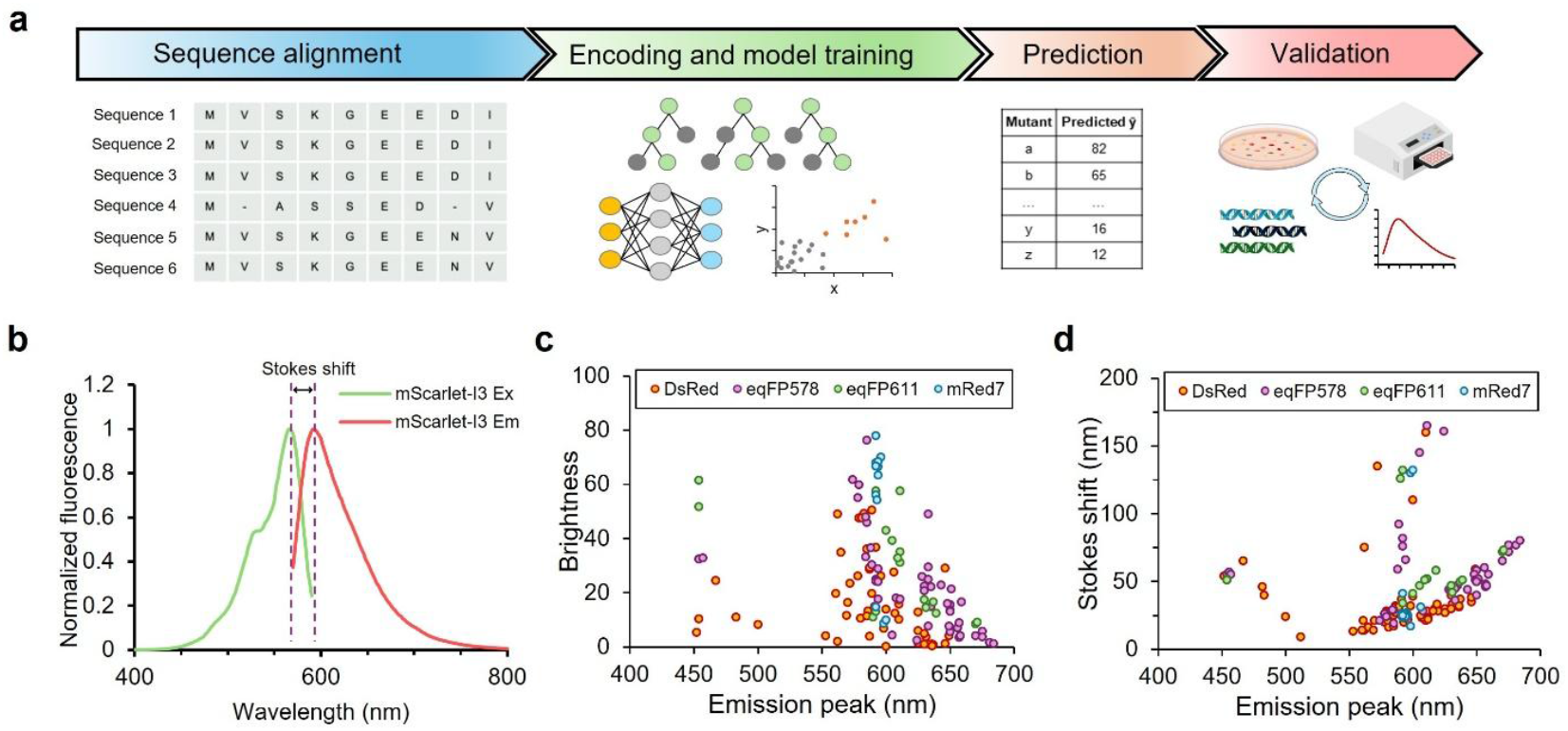
Overview of the dataset and method. (a) Pipeline of the ML-driven approach for predicting and experimentally validating the performance of RFPs in this study. (b) Excitation and emission spectra of mScarlet-I3 with the Stokes shift indicated. (c) Plot of brightness versus emission peak and (d) plot of Stokes shift versus emission peak for RFPs from different lineages used in this study.

## Results and Discussion

### Selection of Target Parameters

Three key photophysical parameters of RFPs, including emission peak, Stokes shift, and molecular brightness, were selected as targets for prediction using ML algorithms. These properties are central to the development of FPs. The emission peak represents the wavelength at which maximum fluorescence is generated. A bathochromic (red) shift is typically desirable because it leads to reduced scattering, absorption, and autofluorescence from biological tissues. Red photons with wavelengths over 600 nm are particularly advantageous for *in vivo* imaging. Stokes shift was calculated as the difference between the emission and excitation peaks of the protein (**Fig. 1b**). RFPs with large Stokes shifts are valuable for single-wavelength excitation and dual-emission microscopy with GFPs and their derived indicators, as their excitation spectra substantially overlap with those of GFPs^30^. Molecular brightness, defined as the product of a protein’s extinction coefficient and quantum yield, is critical for imaging sensitivity. Brighter RFPs provide higher signal-to-noise ratios, especially when fused to low-abundance proteins, and allow imaging at lower excitation intensities, thereby reducing phototoxicity.

### Data Preparation

We compiled existing RFP variants from four lineages, whose ancestral proteins are DsRed, eqFP578, eqFP611, and mRed7, from the Fluorescent Protein Database to generate our dataset.

Tandem-dimer protein sequences were excluded to facilitate sequence alignment, resulting in a total of 154 sequences (**Table S1**). The proteins in the dataset span a broad range of emission peaks, Stokes shifts, and molecular brightness values (**Fig. 1c, d**). Sequences were aligned using the online COBALT^31^, resulting in 251 positions per sequence, including gaps (“–”). Each sequence was then encoded as 251 individual categorical features representing the 20 standard amino acids plus the gap.

### Prediction of Spectral Properties

We first evaluated ML models for predicting emission peak and Stokes shift. The overall process is illustrated in **Fig. 2a**. Amino acid descriptors represent multi-dimensional physicochemical properties of residues, providing an effective way to encode protein sequences. We assessed several common descriptors, including Z-scales^32^, VHSE^33^, T-scales^34^, E-descriptors^35^, and PC scores^36,37^, and compared their performance. Tree-based models, random forest (RF) and XGBoost (XGB), as well as support vector regressor (SVR) were trained with these encodings. The dataset was split 80:20 into training and test sets. Hyperparameters were optimized via Bayesian optimization, and model performance was evaluated on the test set using R^2^ and MAE. Across all metrics, tree-based models outperformed SVR for both emission peak and Stokes shift (**Fig. S1**). Top-ranked models achieved high accuracy, with R^2^ > 0.8 for both properties (**Fig. 2b, c**).

**Fig. 2.**
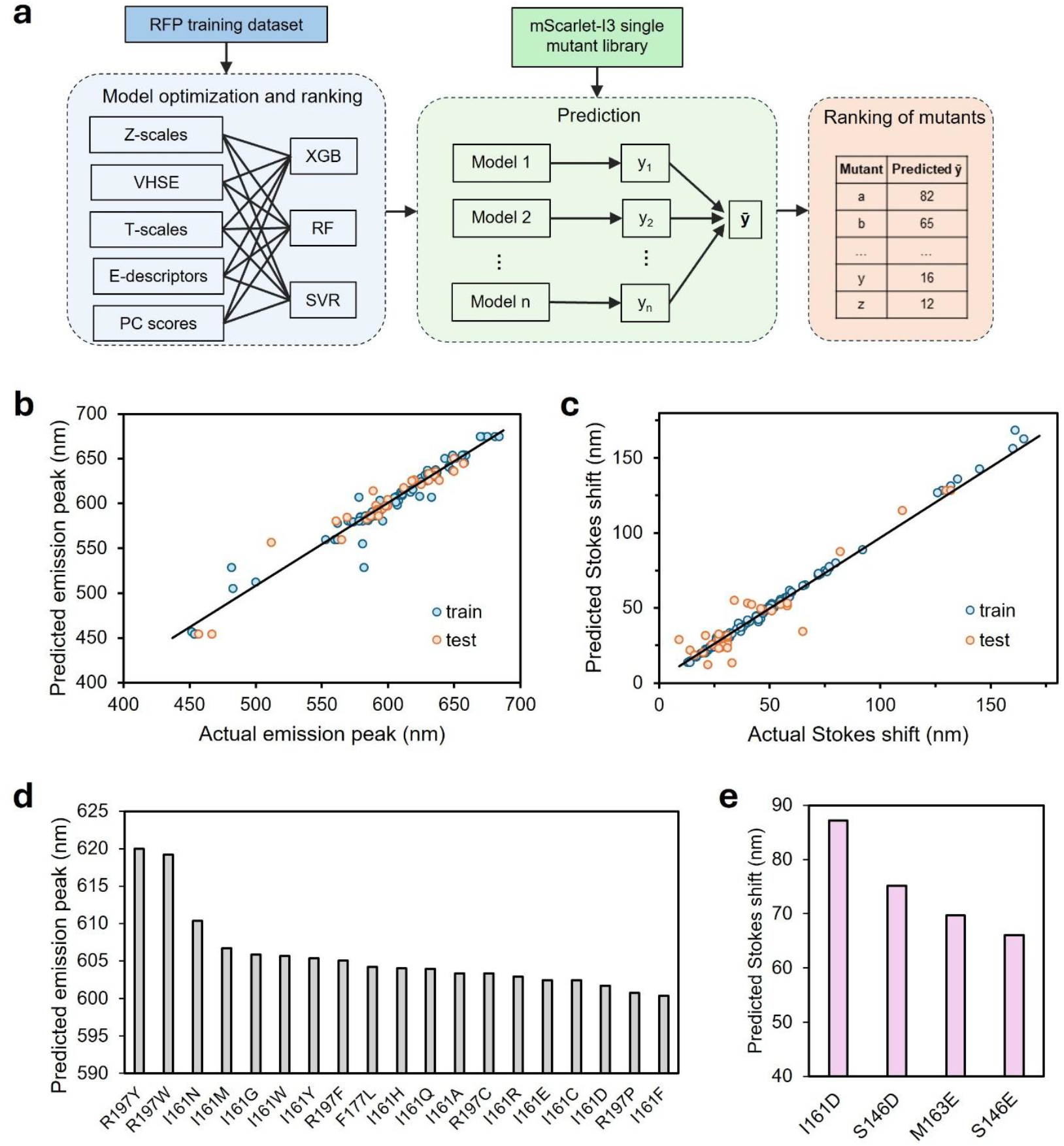
ML regression models for prediction of spectral properties of RFP and mScarlet-I3 mutants. (a) Pipeline for model training and prediction. (b-c) Performance of representative models for predicting the emission peak using random forest with PC scores (b) and Stokes shift using XGBoost with PC scores (c). RFP dataset was used. Solid curves represent linear fitting of the test data points. (d) Top-ranked predicted mScarlet-I3 mutants with red-shifted emission peaks. (e) Top-ranked predicted mScarlet-I3 mutants with increased Stokes shift.

mScarlet-I3 (**Fig. 1b, S2**) is one of the best-performing RFPs, exhibiting ultrafast maturation and high molecular brightness. We next asked whether ML models could predict mutants of mScarlet-I3 with altered spectral properties. Single mutants provide a simple system to evaluate model performance by removing the complexity of multiple interacting mutations. Recent studies also highlight the functional relevance of single mutants of mScarlet-I3 for advanced microscopy^38^. Therefore, we created a single-mutant library by introducing all 20 amino acids at 153 positions in mScarlet-I3, focusing on residues located in β-strands (inward-facing), the central α-helix, and loops. Using this library, we applied the top 4-5 models in an ensemble to predict emission peak and Stokes shift (**Fig. S1**). Mutants were ranked based on predicted values and top ranked mutants were summarized in **Fig. 2d** and **2e**.

For emission peak, mutations at residues R197 and I161 predominantly ranked among the top 20 predicted red-shifted variants. Aromatic residues (Y and W) at R197 were predicted to strongly red-shift emission to over 610 nm, whereas top mutations at I161 included a variety of side chains. For Stokes shift, mutations to D or E at positions I161, S146, and M163 were predicted to produce the largest increase.

### Experimental Validation of Altered Spectral Properties from Prediction

We validated the top ML-predicted mutations using bacterial expression and fluorescence measurements. Deterministic codons were used to generate R197W and R197Y variants. Impressively, R197Y exhibited a pronounced red-shifted emission at 616 nm, consistent with the prediction, albeit with substantially decreased brightness (**Fig. 3a, b**). R197W showed no detectable fluorescence or transmitted color, likely due to improper folding or lack of chromophore formation.

**Fig. 3.**
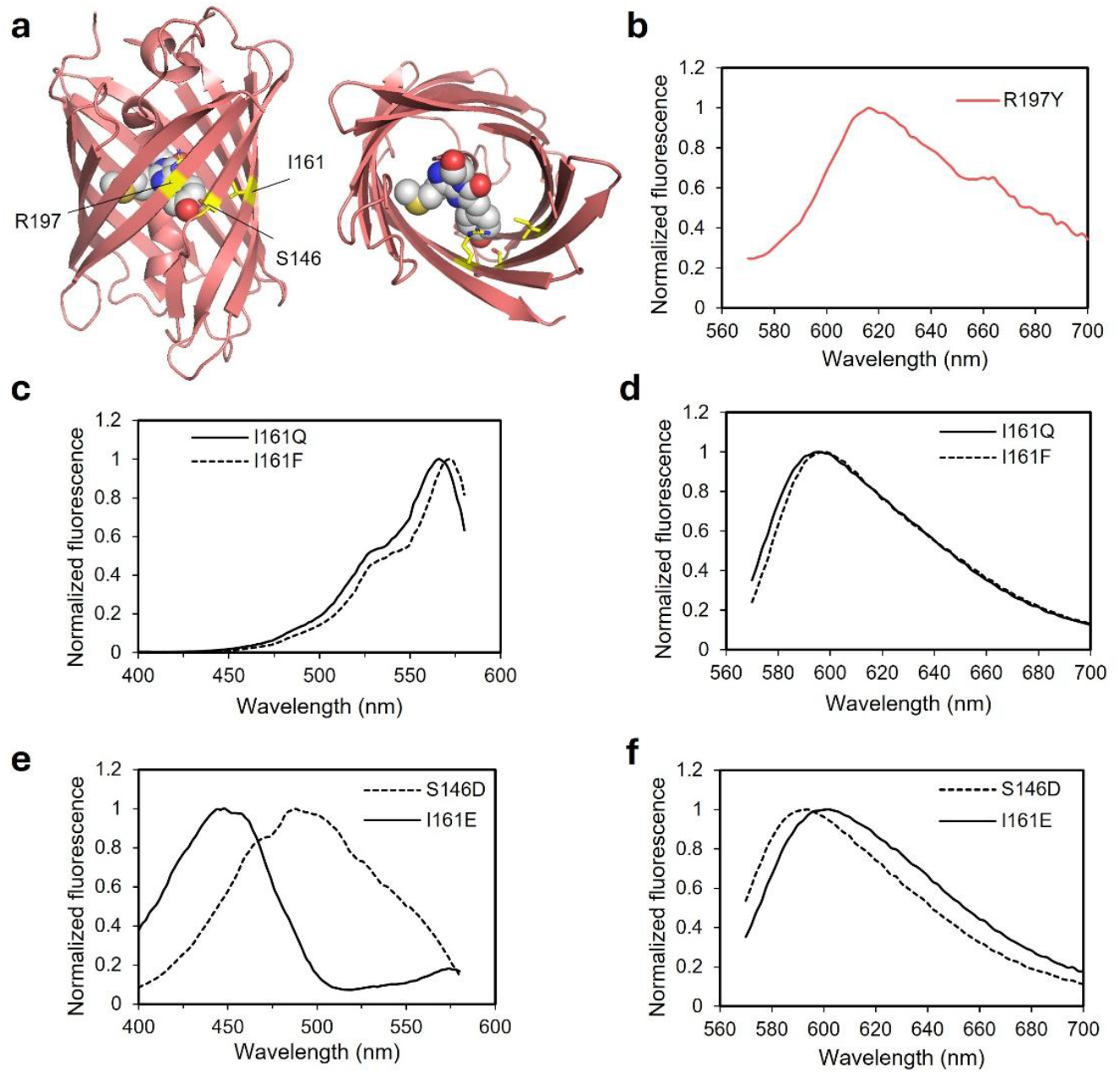
Experimental validation of top-ranked mScarlet-I3 mutants with red-shifted emission and large Stokes shift predicted by ML models. (a) Structure of mScarlet-3 (PDB ID: 7ZCT) with the highest-ranked positions highlighted in yellow. Left: side view. Right: top view. (b) Emission spectrum of mScarlet-I3 R197Y. (c) Excitation and (d) emission spectra of mScarlet-I3 I161Q and I161F. (e) Excitation and (f) emission spectra of mScarlet-I3 S146D and I161E.

Because a range of amino acids were predicted to red-shift emission at I161 (**Fig. 2d**), we employed the degenerate codon NNK to construct a library containing all potential substitutions for screening red-shifted variants. Approximately 300 colonies were screened on agar plates using a fluorescence imager under red fluorescence channel, which effectively covers ten times the library size, ensuring all variants were sampled. Approximately 50 colonies displaying red fluorescence with varying brightness were selected for subculture, lysis, and spectral scanning. This screen identified I161F and I161Q, with emission peaks of 598 nm and 596 nm, respectively (**Fig. 3c, d**), both of which are red-shifted relative to 592 nm for mScarlet-I3.

To validate predicted mutations for increased Stokes shift, degenerate codon GAN were used to introduce D or E at top-ranked positions I161, S146 and M163. Screening revealed that I161E and S146D produced a large Stokes shift, driven by a blue-shifted excitation peak at 448 and 488 nm, respectively (**Fig. 3e, f**). I161E also exhibited red-shifted emission peak at 602 nm, which aligned with the prediction (**Fig. 2d**). S146E, I161D and M163D/E did not substantially change Stokes shift (**Table 1**). Of the two mutations linked to the large Stokes shift phenotype, S146D was previously reported in LSS-mScarlet variants^39,40^, while I161E, to our knowledge, is newly identified as contributing to the large Stokes shift effect in the mScarlet family.

**Table 1.** Spectral properties of mScarlet-I3 mutants in validation of the large Stokes shift phenotype.

| **Mutant** | **Excitation peak**<br>**(nm)** | **Emission peak**<br>**(nm)** | **Stokes shift**<br>**(nm)** |
| --- | --- | --- | --- |
| mScarlet-I3 | 568 | 592 | 24 |
| S146D | 488 | 594 | 106 |
| S146E | 570 | 598 | 28 |
| I161E | 448 | 602 | 154 |
| I161D | 566 | 592 | 26 |
| M163D/E (phenotype 1, 2)<sup>1</sup> | 570, 562 | 598, 596 | 28, 24 |

<sup>1</sup> M163D and M163E were not sequenced following spectral scanning. The specific mutation between the two phenotypes was therefore undetermined.

### Mechanistic Insights into Predicted Spectral Properties

The top-ranked mutations with aromatic side chains (R197W, R197Y, and R197F) in the prediction of red-shifted emission peak are consistent with established structural principles. These substitutions likely promote π-π stacking interactions that lower the energy barrier between the excited and ground states^41^. The models also highlighted I161 as a hotspot, predicting that substitutions to a wide range of amino acids would induce a red shift. Although less obvious from known structural mechanisms, our experiments confirmed that several I161 substitutions (F/Q/E) indeed produced significant red shifts (**Fig 3d, f**), revealing insights that would be difficult to obtain from structural analysis and reasoning alone.

For large Stokes shift, the highest-ranked predictions involved mutations to D or E at positions close to the chromophore’s tyrosine. Introducing negatively charged side chains at these sites can stabilize the phenolate group and facilitate excited-state proton transfer (ESPT), which is the key mechanism underlying large Stokes shift ^42^. We speculate that LSS-mScarlet variants^39,40^ containing S146D in the training set contributed strongly to these predictions, illustrating how the model leveraged subtle patterns in the data.

### Prediction of Molecular Brightness

To predict RFP molecular brightness, we similarly evaluated multiple combinations of amino acid descriptors and ML models and ranked them by performance (**Fig. S3**). The top-ranked models achieved a test R^2^ ∼ 0.6 and MAE ∼ 10, notably lower than the accuracies for emission peak or Stokes shift, indicating that brightness is more difficult to predict from sequence alone using our ML framework. Tree-based algorithms outperformed KNN, SVR, and MLP, so we used the three best RF and three best XGB models as an ensemble to predict the mScarlet-I3 single-mutant library. Mutants were ranked by predicted brightness (**Table S2**). Rather than focusing on individual substitutions, we then ranked mScarlet-I3 positions by the frequency of their appearance among the top 50 predicted mutants (**Fig. 4a**). This approach facilitates experimental validation by enabling the use of degenerate NNK codons to screen the top-ranked positions, providing broader and more efficient coverage than evaluation of each mutation individually.

**Fig. 4.**
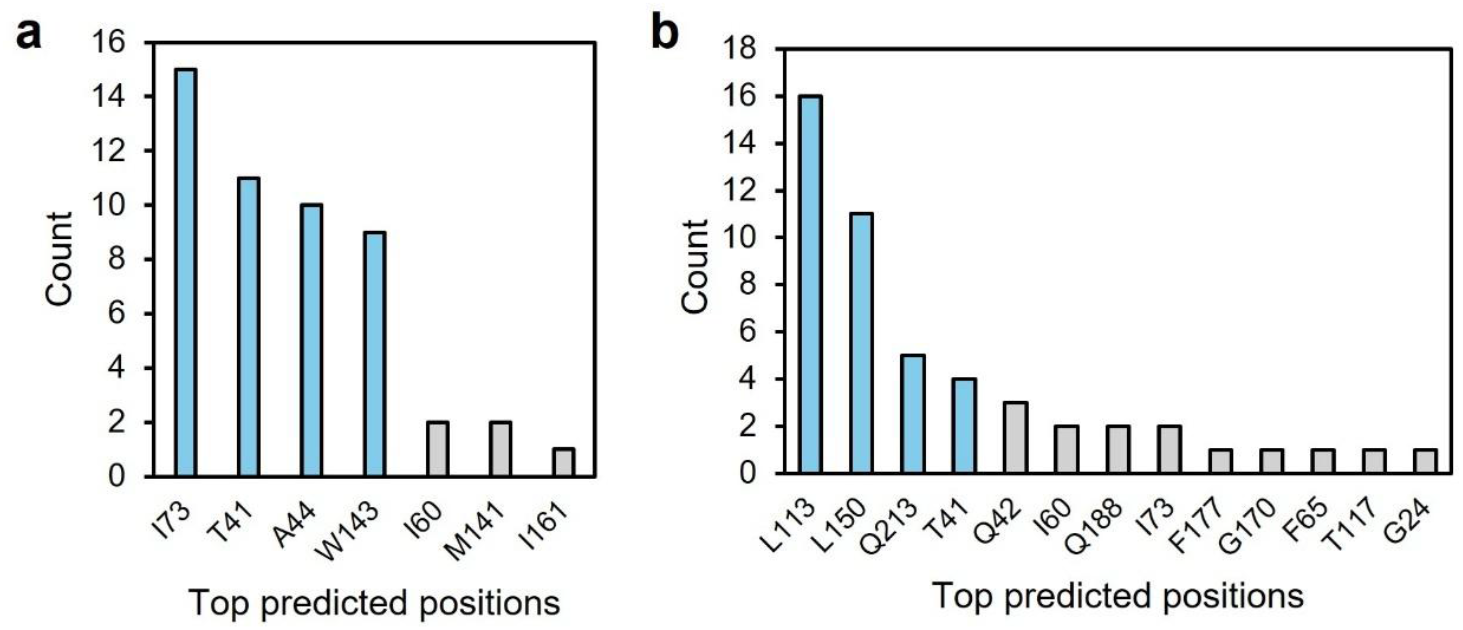
Comparison of ML models for prediction of mScarlet-I3 variant brightness. Counts of positions ranked in the top 50 predicted high brightness using ensemble models with classical amino acid descriptors (a) and optimized AAindex (b). Columns in light blue represent the top positions selected for experimental validation.

Because our ML models using standard amino acid descriptors achieved only moderate accuracy, we next explored alternative strategies to improve performance. A previous study demonstrated that an ensemble ML approach could guide the design of FP-based Ca^2+^ sensors with superior sensitivity^24^. That strategy leveraged a collection of amino acid descriptors (AAindex database^37^) containing 554 one-dimensional physicochemical descriptors. ML models were trained separately with each AAindex descriptor, evaluated on the training set, and the top-performing descriptors were combined with different ML models to build an ensemble for prediction.

Following a similar approach, we used the AAindex database and trained ML models with each of the 554 descriptors on our RFP dataset (**Fig. S4a**). For each ML model (XGB, RF, SVR, and MLP), we selected the five highest-performing AAindex descriptors and combined them to create a five-dimensional representation. Evaluation on the test set confirmed that the best model, XGB, achieved R^2^ > 0.7 and MAE < 8 (**Fig. S4b, c**), an improvement over models using classical amino acid descriptors (**Fig. S3**). Ensemble predictions that combined XGB, RF, SVR, and MLP maintained similar performance. We therefore chose an ensemble excluding MLP for final prediction of the mScarlet-I3 single-mutant library, as it performed slightly better than the ensemble including MLP. Positions were again ranked by the frequency of their mutants appearing among the top 50 predicted for brightness (**Fig. 4b, Table S3**).

### Experimental Validation of Brightness Prediction

We next carried out experimental tests to evaluate the performance of brightness predictions based on both the classical amino acid descriptors and AAindex described above. For screening, we selected the top-ranked positions I73, T41, A44, and W143 from predictions using the classical amino acid descriptors, and L113, L150, and Q213 from predictions using the AAindex (**Fig. 4**). Note that T41 is also among the top 4 positions in the models using the new amino acid descriptors. Libraries of saturation mutagenesis were generated for each of these single positions using the degenerate NNK codon, chosen for its time- and cost-effectiveness (**Fig. 5a**). Each single-position library was plated separately on agar plates with more than 300 colonies to ensure coverage of all 20 possible amino acid substitutions. Colonies were imaged for fluorescence, and the four brightest colonies from each plate were subcultured for plasmid recovery and sequencing. We anticipated that sequencing results would reveal the brightest variants at each site, whether they carried mutations or retained the original residue. Consistent with this expectation, the original amino acid was present among the top four brightest clones at all seven positions (**Fig. 5b**). Notably, A44 and W143 each had all four of the brightest colonies matching the wild-type residue, suggesting that these positions are highly conserved and critical for maintaining brightness. In contrast, more diverse mutations were found among the top candidates at the other positions.

**Fig. 5.**
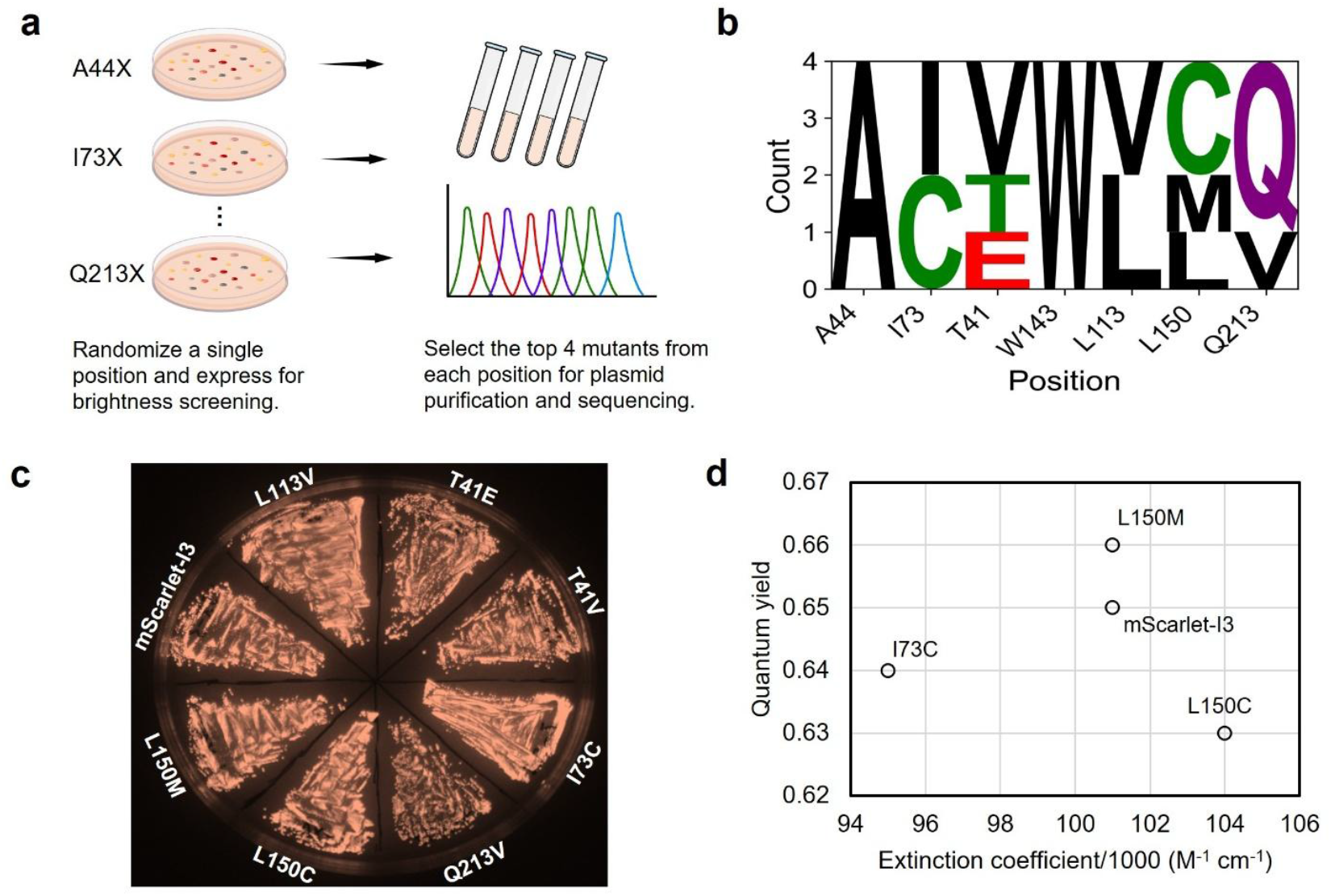
Experimental validation of top-ranked positions from the brightness prediction. (a) Workflow for generating and screening single-position libraries of mScarlet-I3. (b) Sequence-logo plot of the top four mutants selected at each position. (c) Fluorescence images of the selected top mutants expressed in E. coli. (d) Extinction coefficient and quantum yield of selected high brightness variants.

We transformed these top-performing mutants into bacteria and cultured them on the same plate for side-by-side comparison. All of the mutants displayed high and comparable fluorescence intensities, with the exception of Q213V, which was noticeably dimmer (**Fig. 5c**). To accurately quantify brightness, we purified L150M, L150C, and I73C as representatives along with the wild-type mScarlet-I3 and measured their extinction coefficients and quantum yields for comparison. All these variants exhibited comparable brightness, with L150M slightly brighter than the rest (**Fig. 5d, Table 2**). These validations confirmed that models using both amino acid descriptor sets can effectively predict and aid in the discovery of high-brightness variants.

**Table 2.** Characterization of purified mScarlet-I3 mutants.

| **Mutant** | **Extinction coefficient**<br>**(M<sup>-1</sup> cm<sup>-1</sup>)** | **Quantum**<br>**yield** | **Brightness** |
| --- | --- | --- | --- |
| mScarlet-I3 | 101,000 | 0.65 | 65.6 |
| L150C | 104,000 | 0.63 | 65.5 |
| L150M | 101,000 | 0.66 | 66.7 |
| I73C | 95,000 | 0.64 | 60.8 |

### Potential and Limitations of the Models

Our models achieved high accuracy for predicting emission peak and Stokes shift, demonstrating that lightweight AI approaches can capture important aspects of chromophore photophysics. For example, it correctly prioritized mutations that create π-π stacking to red-shift emission and that enable ESPT for large Stokes shift. These are the strategies typically proposed only by experienced fluorescent protein engineers. Moreover, the identification of multiple effective substitutions at I161 for red-shifted emission highlights how ML can uncover non-obvious sequence-function relationships hidden within large sequence datasets.

Compared to the high-accuracy models for spectral properties, the predictions of molecular brightness were only moderately accurate. Brightness depends on both extinction coefficient and quantum yield, which are shaped by complex interactions between the chromophore and multiple surrounding residues. With a small dataset and no explicit structural information, our models inevitably faced limitations. Furthermore, mScarlet-I3 is already among the brightest RFPs known, meaning its brightness is near the upper performance boundary of the dataset. Tree-based models, while outperforming other algorithms, cannot reliably extrapolate beyond the range of the training data. Moreover, designing mutants that surpass state-of-the-art FPs is inherently difficult. For example, while emerging AI approaches have produced mutants that outperform their relatively dim template proteins such as avGFP^17,25^, these improvements have not reached the brightness levels of state-of-the-art GFPs. The brightest FPs to date (e.g., mNeonGreen^3^, StayGold^43^) were solely discovered by exploring natural diversity and then refined through traditional protein engineering rather than computational prediction.

Our results also provide a useful benchmark for evaluating future AI models in protein engineering. One key limitation is the small size of available training datasets, which inherently restricts the predictive accuracy of such models. Pretrained protein language models offer a potential solution to this challenge, as they capture information from vast sequence corpora and can be fine-tuned on smaller experimental datasets to extract relevant sequence-function relationships^44^. However, predicting photophysical properties such as extinction coefficient and quantum yield remains particularly difficult. These properties are highly sensitive to subtle interactions between the chromophore and its surrounding protein environment, and current AI-based structural models have limited ability to accurately capture chromophore-specific photochemistry. Consequently, integrating additional structural information from experimentally determined structures or physics-based simulations could further enhance predictive performance, as demonstrated in previous studies aimed at generating brighter GFP variants^27^. By combining sequence-based ML approaches with structural insights, it may be possible to overcome some of the inherent limitations posed by small datasets and the complex nature of photophysical parameters.

A broader challenge, not unique to our study, is the scarcity of large, balanced datasets for supervised learning in protein engineering. The spectacular success of AI in protein structure prediction was enabled by decades of experimental structure determinations, whereas comparable datasets for fluorescence properties are far smaller and insufficient. Publicly available sequence-function data are biased toward successful or improved proteins, while countless low-performing variants remain unreported, which introduces “survivorship bias”. High-throughput approaches such as flow cytometry-based screening coupled with deep sequencing can help generate genotype-phenotype maps with tens of thousands of data points^45^, but their accuracy and standardization remain areas of investigation.

## Conclusion

In this study, we applied ML models with amino acid descriptor encoding to predict the sequence-function relationships of RFP variants. We demonstrated that lightweight ML models can extract actionable design principles for FPs. Future progress may depend on larger and more representative datasets, integration of structural and photophysical modeling, and continued exploration of hybrid strategies that combine AI with experimental discovery.

## Acknowledgements

The work was supported by NRC AI4D program, Canada Foundation for Innovation (35285) and Ontario Research Fund (35285) to K.P., NSERC Discovery Grant (RGPIN-2025-06922) to Y.Z. and NRC-CRAFT Project Award to K.P. and Y.Z.. We thank Dr. Shengtian Sang for helpful discussion and Victor Sit for technical support.

## Author contributions

Y.Z. conceived the project and collected the training dataset. R.J., A.W., and E.Y.X. performed the experimental validation. J.J., H.C., and Y.Z. conducted the AI-based predictions. K.P. and Y.Z. provided resources and supervised the research. K.P. and Y.Z. wrote the manuscript.

## Supporting Information

Comparison of individual models for predicting emission peaks and Stokes shift. Aligned sequence of mScarlet-I3. Comparison of models using classical amino acid descriptors and AAindex for predicting brightness. List of RFPs used for model training. Top brightness-enhancing mutations predicted by the models. Experimental procedures.

