## Supplementary information for "Predicting and Designing Red Fluorescent Protein Variants Using Sequence-to-Function Machine Learning Models"

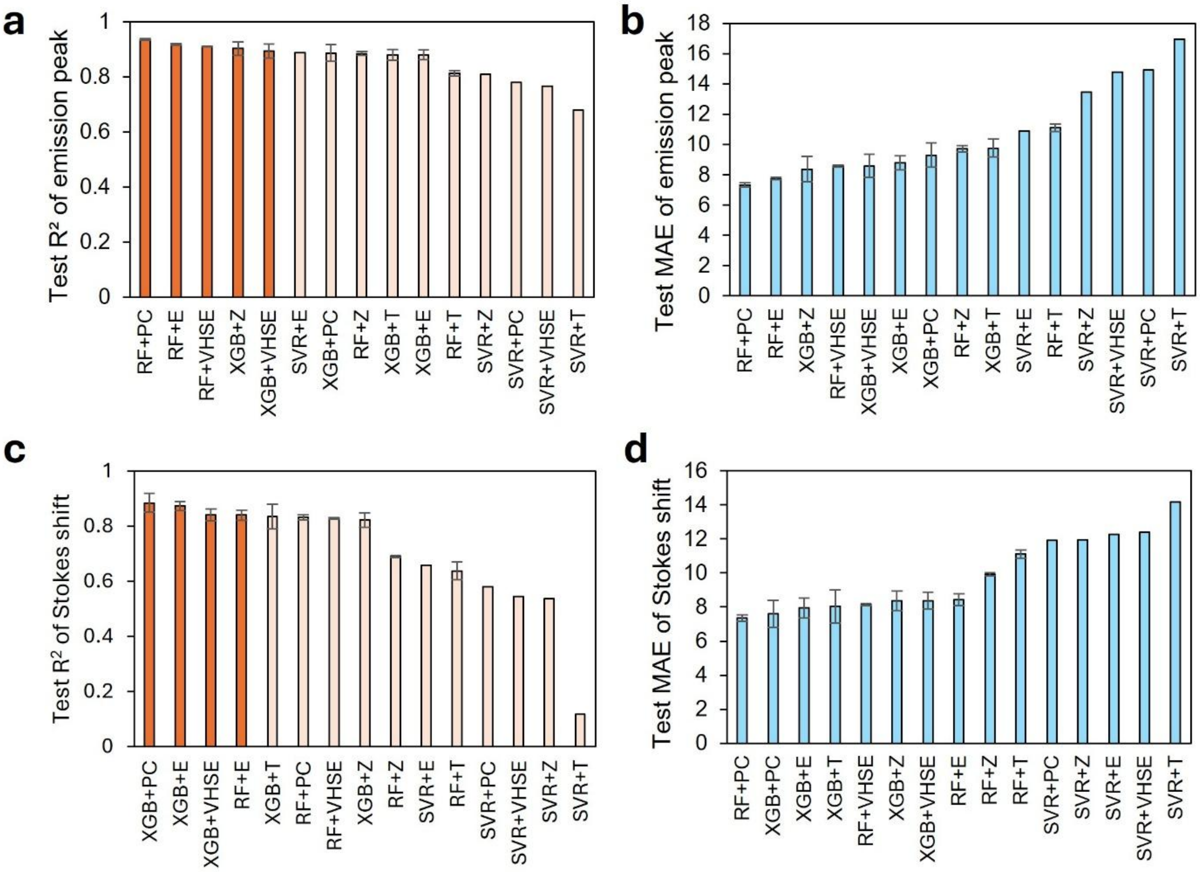


***Fig. S1. Comparison of ML regression models for prediction of spectral properties of RFPs.*** *(a-b) Performance metrics ($R^2$ and MAE) of different ML model and amino acid descriptor combinations for predicting the emission peak. (c-d) Performance metrics ($R^2$ and MAE) of different model and amino acid descriptor combinations for predicting the Stokes shift. RFP dataset was used. Dark-orange columns in (a) and (c) indicate the models used to predict mScarlet-I3 variants. Error bars in (a-d) represent the standard deviation calculated from 10 different random states.*

| | | | | | | | | | | | | | | | | | | | |
|---|---|---|---|---|---|---|---|---|---|---|---|---|---|---|---|---|---|---|---|
| | | | | 5 | | | | | 10 | | | | | 15 | | | | | 20 |
| M | D | S | T | - | E | A | - | - | - | - | V | I | K | E | F | M | R | F | K |
| | | | | 25 | | | | | 30 | | | | | 35 | | | | | 40 |
| V | H | M | E | G | S | M | N | G | H | E | F | E | I | E | G | E | G | E | G |
| | | | | 45 | | | | | 50 | | | | | 55 | | | | | 60 |
| R | P | Y | E | G | T | Q | T | A | K | L | K | V | T | K | G | G | P | L | P |
| | | | | 65 | | | | | 70 | | | | | 75 | | | | | 80 |
| F | S | W | D | I | L | S | P | Q | F | M | Y | G | S | R | A | F | I | K | H |
| | | | | 85 | | | | | 90 | | | | | 95 | | | | | 100 |
| P | A | D | I | P | D | Y | W | K | Q | S | F | P | - | - | - | E | G | F | K |
| | | | | 105 | | | | | 110 | | | | | 115 | | | | | 120 |
| W | E | R | V | M | I | F | E | D | G | G | T | V | S | V | T | Q | D | T | S |
| | | | | 125 | | | | | 130 | | | | | 135 | | | | | 140 |
| L | E | D | G | T | L | I | Y | K | V | K | L | R | G | G | N | F | P | P | D |
| | | | | 145 | | | | | 150 | | | | | 155 | | | | | 160 |
| G | P | V | M | Q | K | R | T | M | G | W | E | A | S | T | E | R | L | Y | P |
| | | | | 165 | | | | | 170 | | | | | 175 | | | | | 180 |
| E | D | V | V | L | K | G | D | I | K | M | A | L | R | L | K | D | G | G | R |
| | | | | 185 | | | | | 190 | | | | | 195 | | | | | 200 |
| Y | L | A | D | F | K | T | T | Y | K | A | K | K | P | - | V | Q | M | - | - |
| | | | | 205 | | | | | 210 | | | | | 215 | | | | | 220 |
| P | G | A | F | N | I | D | R | K | L | D | I | T | S | H | N | E | D | Y | T |
| | | | | 225 | | | | | 230 | | | | | 235 | | | | | 240 |
| V | V | E | Q | Y | E | R | S | V | A | R | H | - | - | - | - | - | S | T | G |
| | | | | 245 | | | | | 250 | | | | | | | | | | |
| - | - | - | - | - | G | S | G | G | - | - | | | | | | | | | |

***Fig. S2.*** *Sequence and residue numbering of mScarlet-I3 after multiple sequence alignment with other RFPs.*

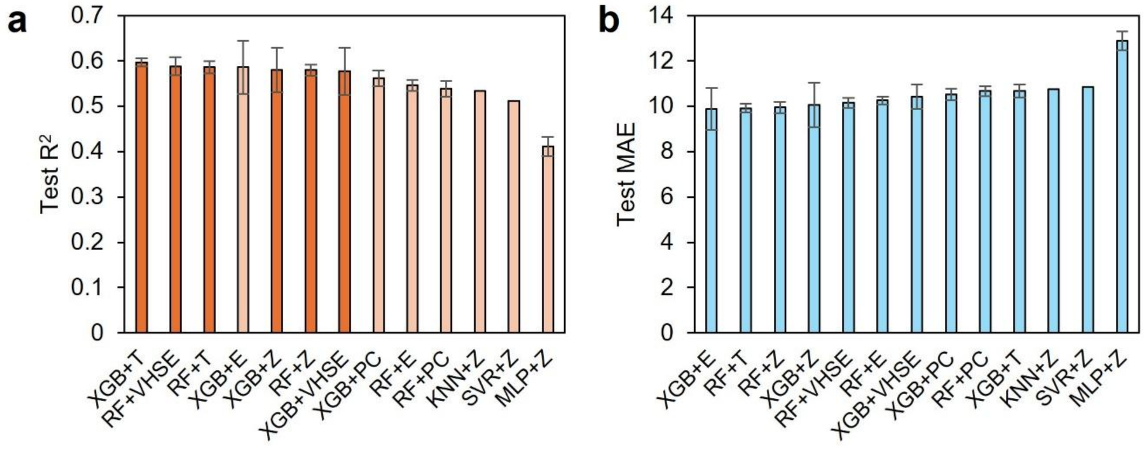


***Fig. S3. Comparison of ML regression models for prediction of RFP brightness using classical amino acid descriptors.*** *(a) $R^2$ and (b) MAE for different combinations of regression models and amino acid descriptors. Dark-orange columns in (a) indicate the models used to predict the mScarlet-I3 variants. Error bars represent the standard deviation calculated from 10 random states.*

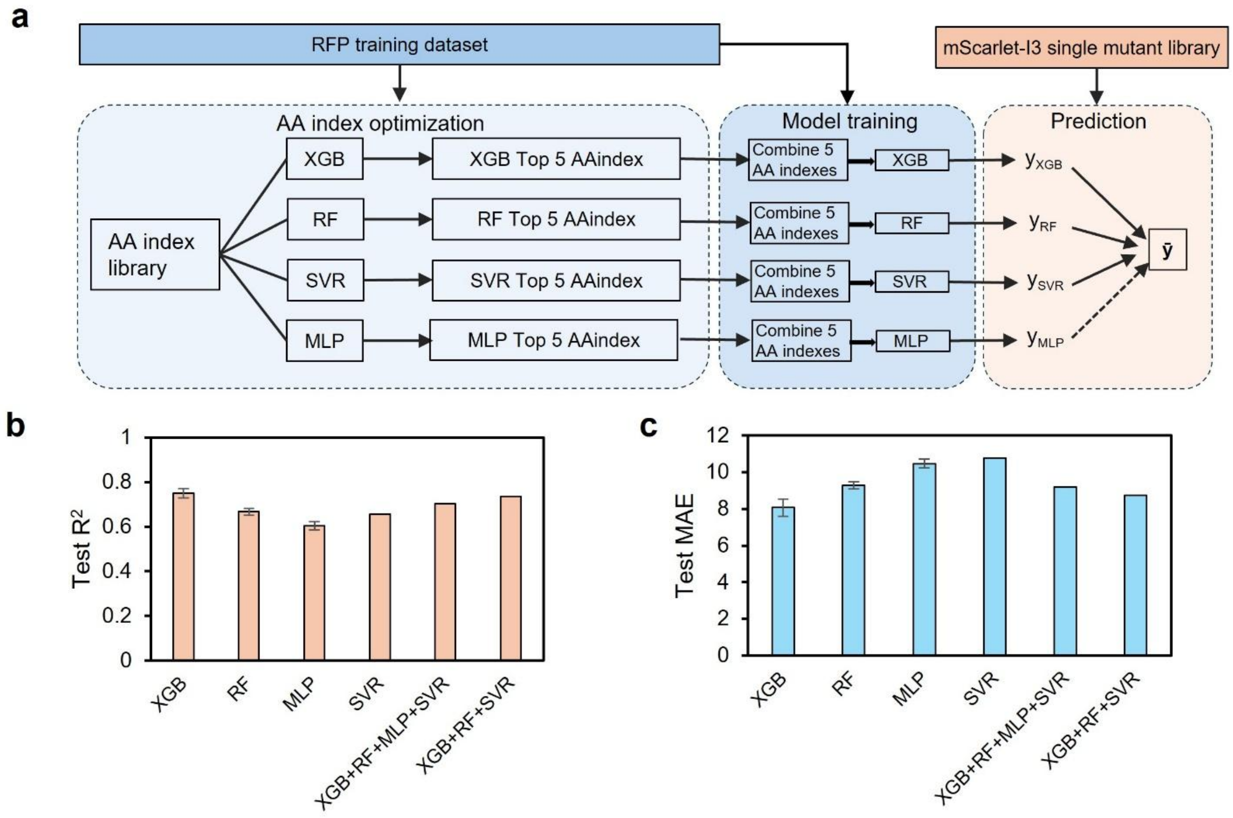


***Fig. S4. Training ML ensemble model using AAindex for RFP brightness prediction.*** *(a) Pipeline for model optimization, training and prediction. (b) R² and (c) MAE for individual model using top 5 AA index and their combinatorial result. Error bars represent the standard deviation calculated from 10 random states.*

**Table S1.** Summary of red fluorescent proteins used in this study.

| | Name | Excitation Peak | Emission Peak | Brightness | Extinction coefficient | Quantum yield | Stokes shift | Family |
|---|---|---|---|---|---|---|---|---|
| 1 | Blue102 | 403 | 454 | 10.26 | 11400 | 0.9 | 51 | DsRed |
| 2 | dimer1 | 551 | 579 | | | | 28 | DsRed |
| 3 | dimer2 | 552 | 579 | 47.61 | 69000 | 0.69 | 27 | DsRed |
| 4 | dis2RFP | 573 | 593 | | | | 20 | DsRed |
| 5 | dis3GFP | 503 | 512 | | | | 9 | DsRed |
| 6 | dsFP483 | 443 | 483 | 10.99 | 23900 | 0.46 | 40 | DsRed |
| 7 | DspR1 | 556 | 582 | | | | 26 | DsRed |
| 8 | DsRed | 558 | 583 | 49.3 | 72500 | 0.68 | 25 | DsRed |
| 9 | DsRed2 | 561 | 587 | 28.6 | 52000 | 0.55 | 26 | DsRed |
| 10 | DsRed-Express | 554 | 586 | 12.64 | 30100 | 0.42 | 32 | DsRed |
| 11 | DsRed-Express2 | 554 | 591 | 14.95 | 35600 | 0.42 | 37 | DsRed |
| 12 | DsRed.M1 | 558 | 587 | 3.82 | 27300 | 0.14 | 29 | DsRed |
| 13 | DsRed-Max | 560 | 589 | 19.68 | 48000 | 0.41 | 29 | DsRed |
| 14 | DsRed-NF | 561 | 600 | 0.06 | 57000 | 0.001 | 39 | DsRed |
| 15 | DsRed.T3 | 560 | 587 | 29.2 | 49500 | 0.59 | 27 | DsRed |
| 16 | DsRed.T4 | 555 | 586 | 13.33 | 30300 | 0.44 | 31 | DsRed |
| 17 | DstC1 | 436 | 482 | | | | 46 | DsRed |
| 18 | dTomato | 554 | 581 | 47.61 | 69000 | 0.69 | 27 | DsRed |
| 19 | E2-Crimson | 611 | 646 | 28.98 | 126000 | 0.23 | 35 | DsRed |
| 20 | E2-Crimson-NF | 583 | | | | | | DsRed |
| 21 | E2-Orange | 540 | 561 | 19.71 | 36500 | 0.54 | 21 | DsRed |
| 22 | E2-Red/Green | 560 | 585 | 36.05 | 53800 | 0.67 | 25 | DsRed |
| 23 | FP586 | 559 | 586 | | | | 27 | DsRed |
| 24 | GdT | 476 | 500 | 8.26 | 59000 | 0.14 | 24 | DsRed |
| 25 | LSSmApple | 490 | 600 | 13.95 | 45000 | 0.31 | 110 | DsRed |
| 26 | LSSmCherry1 | 450 | 610 | 10.15 | 35000 | 0.29 | 160 | DsRed |
| 27 | LSSmOrange | 437 | 572 | 23.4 | 52000 | 0.45 | 135 | DsRed |
| 28 | mApple | 568 | 592 | 36.75 | 75000 | 0.49 | 24 | DsRed |
| 29 | mBanana | 540 | 553 | 4.2 | 6000 | 0.7 | 13 | DsRed |
| 30 | mBlueberry1 | 398 | 452 | 5.28 | 11000 | 0.48 | 54 | DsRed |
| 31 | mBlueberry2 | 402 | 467 | 24.48 | 51000 | 0.48 | 65 | DsRed |
| 32 | mCherry | 587 | 610 | 15.84 | 72000 | 0.22 | 23 | DsRed |
| 33 | mCherry2 | 589 | 610 | 17.47 | 79400 | 0.22 | 21 | DsRed |
| 34 | mCherry_I202T | 587 | 617 | | | 0.04 | 30 | DsRed |
| 35 | mCherry_I202Y | 592 | 620 | | | 0.02 | 28 | DsRed |
| 36 | mCherry_M10L | 583 | 611 | | | | 28 | DsRed |
| 37 | mCherry-XL | 558 | 589 | 50.4 | 72000 | 0.7 | 31 | DsRed |

| | | | | | | | | |
|---|---|---|---|---|---|---|---|---|
| 38 | mGrape1 | 595 | 625 | 1.5 | 50000 | 0.03 | 30 | DsRed |
| 39 | mGrape2 | 605 | 636 | 0.99 | 33000 | 0.03 | 31 | DsRed |
| 40 | mGrape3 | 608 | 646 | 1.2 | 40000 | 0.03 | 38 | DsRed |
| 41 | mHoneydew | 487 | 562 | 2.04 | 17000 | 0.12 | 75 | DsRed |
| 42 | mNectarine | 558 | 578 | 26.1 | 58000 | 0.45 | 20 | DsRed |
| 43 | mOrange | 548 | 562 | 48.99 | 71000 | 0.69 | 14 | DsRed |
| 44 | mOrange2 | 549 | 565 | 34.8 | 58000 | 0.6 | 16 | DsRed |
| 45 | mPlum | 590 | 649 | 4.1 | 41000 | 0.1 | 59 | DsRed |
| 46 | mPlum-E16P | 590 | 630 | | | 0.14 | 40 | DsRed |
| 47 | mRaspberry | 598 | 625 | 12.9 | 86000 | 0.15 | 27 | DsRed |
| 48 | mRFP1 | 584 | 607 | 12.5 | 50000 | 0.25 | 23 | DsRed |
| 49 | mRFP1.1 | 589 | 612 | | | | 23 | DsRed |
| 50 | mRFP1.2 | 590 | 612 | | | | 22 | DsRed |
| 51 | mRFP1_Magenta | 588 | 616 | | | | 28 | DsRed |
| 52 | mRFP1_Pink | 565 | 581 | | | | 16 | DsRed |
| 53 | mRFP1-Q66C | 559 | 580 | 10.49 | 31800 | 0.33 | 21 | DsRed |
| 54 | mRFP1-Q66S | 555 | 569 | 11.52 | 32900 | 0.35 | 14 | DsRed |
| 55 | mRFP1-Q66T | 549 | 570 | 16.38 | 38100 | 0.43 | 21 | DsRed |
| 56 | mRFP1_Violet | 586 | 619 | | | | 33 | DsRed |
| 57 | mRFP1_Yellow | 498 | | | | | | DsRed |
| 58 | mRojoA | 597 | 633 | 0.96 | 48000 | 0.02 | 36 | DsRed |
| 59 | mRojoB | 598 | 631 | 3.66 | 61000 | 0.06 | 33 | DsRed |
| 60 | mRouge | 600 | 637 | 0.86 | 43000 | 0.02 | 37 | DsRed |
| 61 | mSandy2 | 581 | 606 | 27.65 | 79000 | 0.35 | 25 | DsRed |
| 62 | mStrawberry | 574 | 596 | 26.1 | 90000 | 0.29 | 22 | DsRed |
| 63 | mTangerine | 568 | 585 | 11.4 | 38000 | 0.3 | 17 | DsRed |
| 64 | pHuji | 566 | 598 | 6.82 | 31000 | 0.22 | 32 | DsRed |
| 65 | RDSmCherry0.1 | 598 | 625 | 5.96 | 59600 | 0.1 | 27 | DsRed |
| 66 | RDSmCherry0.2 | 600 | 630 | 1.03 | 34400 | 0.03 | 30 | DsRed |
| 67 | RDSmCherry0.5 | 604 | 636 | 0.47 | 23300 | 0.02 | 32 | DsRed |
| 68 | RDSmCherry1 | 600 | 630 | 4.99 | 55400 | 0.09 | 30 | DsRed |
| 69 | sfOrange | 546 | 560 | | | | 14 | DsRed |
| 70 | cgfmKate2 | 584 | 628 | 25.85 | 55000 | 0.47 | 44 | eqFP578 |
| 71 | cgfTagRFP [1] | 556 | 585 | 45.9 | 90000 | 0.51 | 29 | eqFP578 |
| 72 | CyOFP1 | 497 | 589 | 30.4 | 40000 | 0.76 | 92 | eqFP578 |
| 73 | CyRFP1 | 529 | 588 | 36.48 | 48000 | 0.76 | 59 | eqFP578 |
| 74 | dCyOFP2s | 510 | 592 | 24.84 | 36000 | 0.69 | 82 | eqFP578 |
| 75 | dCyRFP2s | 516 | 592 | 24.36 | 42000 | 0.58 | 76 | eqFP578 |
| 76 | eqFP578 | 552 | 578 | 55.08 | 102000 | 0.54 | 26 | eqFP578 |
| 77 | eqFP650 | 592 | 650 | 15.6 | 65000 | 0.24 | 58 | eqFP578 |
| 78 | eqFP670 | 605 | 670 | 4.2 | 70000 | 0.06 | 65 | eqFP578 |

| | | | | | | | | |
|---|---|---|---|---|---|---|---|---|
| 79 | FR-1 | 569 | 594 | 28.87 | 84900 | 0.34 | 25 | eqFP578 |
| 80 | FusionRed | 580 | 608 | 17.95 | 94500 | 0.19 | 28 | eqFP578 |
| 81 | FusionRed-M | 571 | 594 | 24.17 | 71100 | 0.34 | 23 | eqFP578 |
| 82 | FusionRed-MQV | 566 | 585 | 76.32 | 144000 | 0.53 | 19 | eqFP578 |
| 83 | Katushka | 588 | 635 | 22.1 | 65000 | 0.34 | 47 | eqFP578 |
| 84 | Katushka2S | 588 | 633 | 29.48 | 67000 | 0.44 | 45 | eqFP578 |
| 85 | Katushka-9-5 | 588 | 635 | | | | 47 | eqFP578 |
| 86 | LSS-mKate1 | 463 | 624 | 2.5 | 31200 | 0.08 | 161 | eqFP578 |
| 87 | LSS-mKate2 | 460 | 605 | 4.42 | 26000 | 0.17 | 145 | eqFP578 |
| 88 | Maroon0.1 | 610 | 650 | 5.5 | 50000 | 0.11 | 40 | eqFP578 |
| 89 | mBeRFP | 446 | 611 | 17.55 | 65000 | 0.27 | 165 | eqFP578 |
| 90 | mCardinal | 604 | 659 | 16.53 | 87000 | 0.19 | 55 | eqFP578 |
| 91 | mCarmine | 603 | 675 | 5.81 | 83000 | 0.07 | 72 | eqFP578 |
| 92 | mCyRFP1 | 528 | 594 | 17.55 | 27000 | 0.65 | 66 | eqFP578 |
| 93 | mKate | 588 | 635 | 14.85 | 45000 | 0.33 | 47 | eqFP578 |
| 94 | mKate2 | 588 | 633 | 25 | 62500 | 0.4 | 45 | eqFP578 |
| 95 | mKate M41G S158C | 593 | 648 | 16.06 | 73000 | 0.22 | 55 | eqFP578 |
| 96 | mKate S158A | 585 | 630 | 22.5 | 75000 | 0.3 | 45 | eqFP578 |
| 97 | mKate S158C | 586 | 630 | 20.79 | 63000 | 0.33 | 44 | eqFP578 |
| 98 | mKelly1 | 596 | 656 | 7.04 | 44000 | 0.16 | 60 | eqFP578 |
| 99 | mKelly2 | 598 | 649 | 7.74 | 43000 | 0.18 | 51 | eqFP578 |
| 100 | mMaroon1 | 609 | 657 | 8.8 | 80000 | 0.11 | 48 | eqFP578 |
| 101 | mNeptune | 600 | 650 | 13.4 | 67000 | 0.2 | 50 | eqFP578 |
| 102 | mNeptune2 | 599 | 651 | 21.36 | 89000 | 0.24 | 52 | eqFP578 |
| 103 | mNeptune2.5 | 599 | 643 | 22.8 | 95000 | 0.24 | 44 | eqFP578 |
| 104 | mNeptune681 | 604 | 681 | 1.52 | 38000 | 0.04 | 77 | eqFP578 |
| 105 | mNeptune684 | 604 | 684 | 1.17 | 39000 | 0.03 | 80 | eqFP578 |
| 106 | mStable | 597 | 633 | 7.65 | 45000 | 0.17 | 36 | eqFP578 |
| 107 | mTagBFP2 | 399 | 454 | 32.38 | 50600 | 0.64 | 55 | eqFP578 |
| 108 | Neptune | 600 | 650 | 12.96 | 72000 | 0.18 | 50 | eqFP578 |
| 109 | secBFP2 | 399 | 456 | | | | 57 | eqFP578 |
| 110 | super-TagRFP | 555 | 579 | 59.89 | 113000 | 0.53 | 24 | eqFP578 |
| 111 | TagBFP | 402 | 457 | 32.76 | 52000 | 0.63 | 55 | eqFP578 |
| 112 | TagRFP | 555 | 584 | 48 | 100000 | 0.48 | 29 | eqFP578 |
| 113 | TagRFP657 | 611 | 657 | 3.4 | 34000 | 0.1 | 46 | eqFP578 |
| 114 | TagRFP658 | 611 | 658 | 4.12 | 41200 | 0.1 | 47 | eqFP578 |
| 115 | TagRFP675 | 598 | 675 | 3.68 | 46000 | 0.08 | 77 | eqFP578 |
| 116 | TagRFP-T | 555 | 584 | 33.21 | 81000 | 0.41 | 29 | eqFP578 |
| 117 | tdKatushka2 | 588 | 633 | 49.02 | 132500 | 0.37 | 45 | eqFP578 |
| 118 | TurboRFP | 553 | 574 | 61.64 | 92000 | 0.67 | 21 | eqFP578 |
| 119 | CRISPRed2s | 464 | 590 | 10.96 | 28700 | 0.382 | 126 | eqFP611 |

| | | | | | | | | |
|---|---|---|---|---|---|---|---|---|
| 120 | d-RFP618 | 560 | 618 | | | 0.35 | 58 | eqFP611 |
| 121 | Electra1 | 402 | 454 | 51.62 | 69750 | 0.74 | 52 | eqFP611 |
| 122 | Electra2 | 403 | 454 | 61.48 | 80900 | 0.76 | 51 | eqFP611 |
| 123 | eqFP611 | 559 | 611 | 35.1 | 78000 | 0.45 | 52 | eqFP611 |
| 124 | eqFP611 V124T | 559 | 611 | 31.08 | 74000 | 0.42 | 52 | eqFP611 |
| 125 | mCRISPRed | 460 | 592 | 13.11 | 28500 | 0.46 | 132 | eqFP611 |
| 126 | mGarnet | 598 | 670 | 8.55 | 95000 | 0.09 | 72 | eqFP611 |
| 127 | mGarnet2 | 598 | 671 | 9.13 | 105000 | 0.087 | 73 | eqFP611 |
| 128 | mRuby | 558 | 605 | 39.2 | 112000 | 0.35 | 47 | eqFP611 |
| 129 | mRuby2 | 559 | 600 | 42.94 | 113000 | 0.38 | 41 | eqFP611 |
| 130 | mRuby3 | 558 | 592 | 57.6 | 128000 | 0.45 | 34 | eqFP611 |
| 131 | RFP611 | 559 | 611 | 57.6 | 120000 | 0.48 | 52 | eqFP611 |
| 132 | RFP618 | 560 | 618 | | | 0.35 | 58 | eqFP611 |
| 133 | RFP630 | 583 | 630 | 17.5 | 50000 | 0.35 | 47 | eqFP611 |
| 134 | RFP637 | 587 | 637 | 16.56 | 72000 | 0.23 | 50 | eqFP611 |
| 135 | RFP639 | 588 | 639 | 12.42 | 69000 | 0.18 | 51 | eqFP611 |
| 136 | td-RFP611 | 558 | 609 | 32.9 | 70000 | 0.47 | 51 | eqFP611 |
| 137 | td-RFP639 | 589 | 631 | 14.46 | 90400 | 0.16 | 42 | eqFP611 |
| 138 | LSSmScarlet | 470 | 598 | 12.68 | 30200 | 0.42 | 128 | mRed7 |
| 139 | LSSmScarlet2 | 470 | 600 | 8.7 | 30000 | 0.29 | 130 | mRed7 |
| 140 | LSSmScarlet3 | 466 | 598 | 9.83 | 27300 | 0.36 | 132 | mRed7 |
| 141 | mRed7 | 589 | 606 | | | | 17 | mRed7 |
| 142 | mRed7Q1 | 560 | 591 | | | | 31 | mRed7 |
| 143 | mRed7Q1S1 | 569 | 595 | | | | 26 | mRed7 |
| 144 | mRed7Q1S1BM | 569 | 596 | | | | 27 | mRed7 |
| 145 | mScarlet | 569 | 594 | 70 | 100000 | 0.7 | 25 | mRed7 |
| 146 | mScarlet-220A | 568 | 594 | 63.46 | 95000 | 0.668 | 26 | mRed7 |
| 147 | mScarlet-2A | 568 | 592 | 66.44 | 97000 | 0.685 | 24 | mRed7 |
| 148 | mScarlet-2A-84W | 568 | 592 | 67.9 | 97000 | 0.7 | 24 | mRed7 |
| 149 | mScarlet3 | 569 | 592 | 78 | 104000 | 0.75 | 23 | mRed7 |
| 150 | mScarlet-H | 551 | 592 | 14.8 | 74000 | 0.2 | 41 | mRed7 |
| 151 | mScarlet-I | 569 | 593 | 56.16 | 104000 | 0.54 | 24 | mRed7 |
| 152 | mScarlet-I-220A | 568 | 594 | 54.3 | 100000 | 0.543 | 26 | mRed7 |
| 153 | mScarlet-I3 | 568 | 592 | 68.25 | 105000 | 0.65 | 24 | mRed7 |
| 154 | mScarlet-I3-NCwt | 567 | 592 | 66.61 | 102000 | 0.653 | 25 | mRed7 |

[1] The parameters for cgfTagRFP in the Fluorescent Protein Database (https://www.fpbase.org/) may have been updated since our analysis, as they differ slightly from the values shown here. Our model training was conducted using the values provided in this table.

**Table S2.** Top 50 mutations of mScarlet-I3 ranked by the ensemble model for brightness prediction using classical amino acid descriptors in Fig.4a.

| Rank | Mutation[1] | Rank | Mutation | Rank | Mutation | Rank | Mutation | Rank | Mutation |
|---|---|---|---|---|---|---|---|---|---|
| 1 | I73T | 11 | A44M | 21 | T41H | 31 | A44Y | 41 | M141R |
| 2 | W143S | 12 | I73N | 22 | I73R | 32 | A44F | 42 | I73Y |
| 3 | W143A | 13 | W143C | 23 | W143H | 33 | A44P | 43 | T41V |
| 4 | I73S | 14 | A44V | 24 | T41Y | 34 | I73H | 44 | T41I |
| 5 | W143T | 15 | A44I | 25 | I73K | 35 | A44W | 45 | A44N |
| 6 | W143G | 16 | A44L | 26 | T41F | 36 | W143N | 46 | M141E |
| 7 | I73G | 17 | I161V | 27 | W143D | 37 | W143E | 47 | I60G |
| 8 | I73C | 18 | I73P | 28 | T41E | 38 | T41M | 48 | T41N |
| 9 | I73A | 19 | I73E | 29 | I73V | 39 | T41Q | 49 | A44D |
| 10 | I73D | 20 | T41W | 30 | I73Q | 40 | I60V | 50 | T41L |

[1] Residue numbers correspond to the original mScarlet-I3 sequence prior to multiple sequence alignment.

**Table S3.** Top 50 mutations of mScarlet-I3 ranked by the ensemble model for brightness prediction using AAindex in Fig.4b.

| Rank | Mutation[1] | Rank | Mutation | Rank | Mutation | Rank | Mutation | Rank | Mutation |
|---|---|---|---|---|---|---|---|---|---|
| 1 | L113W | 11 | L150W | 21 | Q42C | 31 | L113T | 41 | G170C |
| 2 | L113C | 12 | L113P | 22 | L150P | 32 | Q213S | 42 | T41M |
| 3 | L113H | 13 | L150H | 23 | L150Y | 33 | L150Q | 43 | F65W |
| 4 | L113M | 14 | L113D | 24 | L150N | 34 | Q188M | 44 | I73N |
| 5 | L113Y | 15 | I60L | 25 | T41W | 35 | T41H | 45 | L150S |
| 6 | L150C | 16 | L113I | 26 | L150F | 36 | L113A | 46 | Q213V |
| 7 | L113F | 17 | L150M | 27 | T41C | 37 | Q42H | 47 | I73T |
| 8 | L113N | 18 | Q213E | 28 | Q42W | 38 | Q213K | 48 | T117M |
| 9 | Q213L | 19 | Q188W | 29 | L150D | 39 | I60E | 49 | G24C |
| 10 | L113Q | 20 | F177M | 30 | L113R | 40 | L113G | 50 | L113S |

[1] Residue numbers correspond to the original mScarlet-I3 sequence prior to multiple sequence alignment.

### Experimental Section

#### Data Collection, Sequence Alignment, and Processing

FP sequences and their photophysical properties were obtained from https://www.fpbase.org/. RFP lineages derived from DsRed, eqFP578, eqFP611, and mRed7 were selected for model training, yielding 154 unique sequences. Tandem-dimer proteins (GGvT, RRvT, tdTomato, and tdimer2(12)) were excluded because their dimeric sequences cannot be reliably aligned with the predominantly monomeric proteins in the dataset.

Multiple sequence alignment of the RFPs was performed with Constraint-based Multiple Alignment Tool, COBALT (https://www.ncbi.nlm.nih.gov/tools/cobalt/re_cobalt.cgi). The aligned sequences were encoded as 251 discrete features, each representing a single amino acid position with 20 standard residue symbols plus a "–" to denote gaps. Target variables for model training included molecular brightness (extinction coefficient × quantum yield), emission peak wavelength, and Stokes shift (emission minus excitation peak). Variants lacking any of these target measurements were removed from the training dataset.

#### Model Training and Optimization.

ML models were implemented in Python 3.11.7 using scikit-learn and XGBoost (XGB) within Jupyter Notebook. Datasets were split 80:20 into training and test sets. For models using amino acid descriptors, features were encoded as dictionaries and gap positions ("–") were assigned zeros. The final feature space size equaled 251 positions multiplied by the dimensionality of the chosen descriptor.

For each combination of algorithm and descriptor, hyperparameters were tuned either manually or via Bayesian optimization (3-fold cross-validation, scoring = "neg_mean_absolute_error"). The best-scoring hyperparameters were used for model training, and performance was evaluated as the mean test $R^2$ and mean absolute error (MAE) across ten random seeds. The top-performing algorithm-descriptor combinations, based on $R^2$, were selected for the final ensemble predictions. $R^2$ and MAE is calculated as:

$$R^2 = 1 - \frac{\sum_{i=1}^{n}(y_i - \hat{y}_i)^2}{\sum_{i=1}^{n}(y_i - \bar{y}_i)^2}$$

$$MAE = \frac{1}{n}\sum_{i=1}^{n}|y_i - \hat{y}_i|$$

where $n$= number of samples, $y_i$= actual value, $\hat{y}_i$= predicted value, and $\bar{y}$= mean of the actual values.

For models employing one-dimensional AAindex descriptors, the dataset was again split at 80:20 for train and test. XGB, random-forest (RF) regression, support-vector regression (SVR), and multilayer perceptron (MLP) models were trained on each individual AAindex encoding, and the average test $R^2$ across five random seeds was recorded. The top five AAindex descriptors with the highest $R^2$ for each model were combined into a five-dimensional representation and used for training and final ensemble predictions.

**Prediction of mScarlet-I3 Variants**

To design a single-mutant library of mScarlet-I3, the crystal structure was inspected and each residue categorized as in (inward-facing β-strand), out (outward-facing β-strand), helix (central α-helix), loop (inter-strand linker), or termini (N- or C-terminus). Positions labeled in, helix, or loop (153 total) were selected for mutagenesis. Each site was substituted with the remaining 19 amino acids, generating 153 × 19 = 2,907 single-mutant sequences; however, the list used for prediction contains 153 × 20 = 3,060 entries, as it also includes the unmutated wild-type residue at each position. These sequences were encoded with the optimized amino acid descriptors and input to the trained models to predict emission peak, Stokes shift, and molecular brightness. The ensemble predictions from all models were averaged to generate the final values, which were subsequently ranked based on the target property.

**Experimental Validation**

To construct mScarlet-I3 mutants, overlap PCR was performed using primers carrying the desired mutations with Q5® High-Fidelity 2X Master Mix (NEB). Saturation mutagenesis at each position was achieved using the degenerate codon NNK, covering all 20 amino acids. PCR products were purified by agarose gel electrophoresis and cloned into a pBAD expression vector using NEBuilder® HiFi DNA Assembly. The assembled constructs were transformed into DH10B electrocompetent cells via electroporation and plated on LB-agar supplemented with 100 µg/mL ampicillin and 0.2% arabinose. Plates were incubated overnight at 37 °C.

To screen for red-shifted emission, colonies expressing mScarlet-I3 variants were imaged using a gel imager. Selected colonies were sub-cultured in 4 mL LB medium containing 100 µg/mL ampicillin and 0.2% arabinose in a shaker incubator at 37 °C. The next day, bacteria were harvested by centrifugation, washed with PBS, and lysed using B-PER™ II Bacterial Protein Extraction Reagent (Thermo Fisher Scientific). Lysates were diluted in PBS for fluorescence measurements on a BioTek Synergy Neo2 plate reader (Agilent). Mutants exhibiting red-shifted emission peaks were subjected to plasmid purification and Sanger sequencing.

To screen for brightness, colonies were imaged using a ChemiDoc imager (Bio-Rad). The brightest colonies from each plate were selected for liquid culture, plasmid purification, and sequencing as described above. Selected mutants were expressed and purified using Ni-NTA affinity chromatography, followed by buffer exchange with Amicon ultrafiltration tubes, as previously described. Purified proteins were characterized for molecular brightness. Extinction coefficients were determined by measuring protein absorbance in 0.1 M NaOH (denatured) and PBS (intact) at equal concentrations. The extinction coefficient of the intact protein was calculated by comparing its absorbance to that of the denatured protein at 460 nm, assuming an extinction coefficient of 44,000 $M^{-1} \cdot cm^{-1}$ for the denatured form. Quantum yields were measured using mScarlet-I3 (QY = 0.65) as a reference. Absorbance and fluorescence of each mutant were recorded at different concentrations, and integrated fluorescence was plotted versus absorbance. Slopes from linear fits were compared to the reference to calculate the quantum yield of each mutant.
